# APPLICATION OF PETRI NETS METHODOLOGY FOR THE EXAMINATION OF THE BIO(PHYSICO)CHEMICAL PARAMETERS OF MITOCHONDRIA FUNCTIONING

**DOI:** 10.1101/2020.01.22.915074

**Authors:** Hanna V. Danylovych, Alexander Yu. Chunikhin, Yuriy V. Danylovych, Sergiy O. Kosterin

**Author notes:** corresponding author: Danylovych Hanna.

## Abstract

A mathematical model is developed using Petri net methodology that is able to predict simultaneous changes in bio(physical)chemical parameters of mitochondria, namely hydrodynamic diameter, fluorescence of NADH, DCF fluorescence signal depending on the time of action and concentration of sodium azide and predict these changes for unknown NaN_3_ concentration. The model is created for real experimental conditions, it combines functional changes of mitochondria with a structural representation of learned processes.

## INTRODUCTION

Biochemistry and molecular physiology of mitochondria, which combines the processes of oxygen consumption, oxidative phosphorylation, catabolism of lipids, biosynthesis of heme, maintenance of Ca^2+^ homeostasis, production of reactive oxygen and nitrogen species, apoptosis, etc. are the priority directions of modern biological science [1–3]. Further elucidation of the electronic transport chain functioning mechanisms and ways of breaking mitochondrial bioenergetics is important for understanding the causes and effects of mitochondrial dysfunction, which is the basis of the smooth muscles contractile function pathology. In particular, the generation of ATP and the functioning of low-affinity and high-capasitive Ca^2+^ uniporter significantly affects the path of Ca^2+^ signaling in myocytes and is an important factor in reducing the concentration of this cation in the myoplasm after the Ca^2+^ transient [4–5]. This also applies to the mitochondria of the uterus smooth muscle, where the electrochemical gradient dissipation of the internal membrane leads to a generalized increase in the myoplasm concentration of Ca^2+^ [6]. Inhibitors of some complexes of the respiratory chain are widely used in order to study thise.

Biomarkers of mitochondria functional activity are parameters such as the endogenous fluorescence signal from adenine nucleotides (NADH), the volume of mitochondria (hydrodynamic diameter), the intensity of reactive oxygen species production (DCF fluorescence), the efficiency of the Ca^2+^ accumulation, etc [4, 7–11]. The possibility of simultaneous simulation of these processes is important for understanding the functioning of mitochondria as a holistic system and will enable to predict the consequences of the violation of certain electronic transport chain components for bioenergy, Ca^2+^ homeostasis and programmed cell death. In particular, it seems advisable to construct a model based on data from the known inhibitor of the *IV* complex of the respiratory chain and the "indirect NO donor " [12] sodium azide action on the bio(physical)chemical parameters of the mitochondria. The actuality of this work is due to the widespread use of NaN_3_ as a tool for inhibiting oxidative phosphorylation and transport functions of mitochondria, as well as the possibility of endogenous synthesis of NO in these subcellular structures.

We are developing a simulation model that links changes in the functioning of the electronic transport chain, the production of reactive oxygen species, the endogenous fluorescence of adenine nucleotides, and mitochondria swelling for the formalization and generalization of experimental data, in order to carry out the predictive function, and also to find the correspondence between the theoretical predictions and real results. We use the methodology of hybrid functional Petri nets as a modeling tool. The advantages of hybrid functional Petri nets as a modeling method include the following [13]: (1) capability to structurally represent the states of the modeled system and the processes occurring in the system; (2) quantitative modeling of three types of states and processes simultaneously, namely discrete, continuous, and associative (forming); (3) possibility to consider the activating, inhibiting, and catalytic effects by the means of a special type of bonds.

The aim of the paper was to use Petri nets to create an imitation model for simultaneous changes in the fluorescence of endogenous NADH, the characteristic size of mitochondria and the generation of reactive oxygen species in real experimental conditions, which would combine functional changes with the structural representation of these processes.

## MATERIALS AND METHODS

Experiments were performed on white wild-type nonpregnant rats weighing 150-180 g. All manipulations with animals were carried out according to the European Convention for the Protection of Vertebrate Animals used for Experimental and other Scientific Purposes, and the Law of Ukraine “On protection of animals from cruelty”. Rats were anesthetized by diethyl ether inhalation and decapitated.

### Isolation of mitochondria from the smooth muscle of the uterus (myometrium) of non-pregnant rats

The mitochondrial fraction was isolated from the myometrium of non-pregnant rats using differentiation centrifugation, as described by Kosterin and coworkers [14]. For the duration of the experiment, the isolated mitochondrial fraction was kept on ice. The protein content of the mitochondrial fraction was determined by a standard procedure by Bradford [15]. The total protein content in the mitochondrial fraction was 2 mg/mL.

### Registration of DCF-fluorescence in mitochondria

The loading of mitochondria by reactive oxygen species sensitive DCF-DA fluorescence probe at a concentration of 25 µM was performed in a medium containing 10 mM Hepes (pH 7.4, 37°C), 250 mM sucrose, 0.1% bovine serum albumin for 30 min at 24°C. The Pluronic F-127 dye was added (0.02%) to improve the loading process. DCF-fluorescense in isolated mitochondria was studied using the flow cytometry method on the COULTER EPICS XL^™^ (Beckman Coulter, USA) equipped with an argon laser (λ_ex_ = 488 nm, λ_fl_ = 525 nm (Fl1 channel). The reaction was initiated by adding aliquots (20 µl) of 5 mM pyruvate + 5 mM succinate. Incubation time 35 min. Sodium azide were added for 15 min. The protein content in the aliquot of the mitochondria fraction was 20-25 µg.

### Estimation of mitochondrial hydrodynamic diameter

To assess changes in the mitochondrial volume, we used the Dynamic light scattering method, which allowed us to determine their sizes (average hydrodynamic diameter). The volume of the particles in suspension was analyzed using the correlation spectrometer ZetaSizer-3 (Malvern Instruments, UK) equipped with He-Ne laser LGN-111 (P = 25 mW, λ = 633 nm). Its operation principle is based on the analysis of time-dependent fluctuations in the scattering intensity when a laser ray passes through a medium with the mitochondria. The temporal intensity changes are converted into the mean translational diffusion coefficient (D) [16]. The translational diffusion coefficient is related to the duration of the correlation τ_c_ with the following ratio:

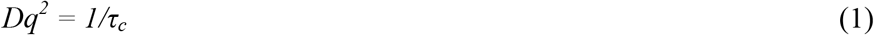

The wave vector of the concentration fluctuations (*q*) is described by the following expression:

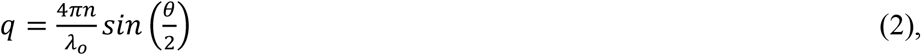

where *n* is the refractive index of the medium (liquid), λ_0_ is the wavelength, and θ is the scattering angle.

Using the Stokes–Einstein equation that connects the particle size, the translational diffusion coefficient, and the viscosity of the liquid, we could calculate the size (diameter) d(H) of the spherical particles as follows [16]:

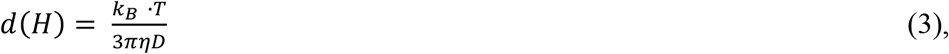

where *k*_*B*_ is the Boltzmann constant; *T* is the absolute temperature, K; η is the shear viscosity of the medium in which the particles are suspended; and *D* is the translational diffusion coefficient.

The recording and statistical processing of the changes in the scattering intensity in the mitochondria water suspension (*n* = 1.33) were performed 10 times for 10 min at +22°C, at a scattering angle of 90°. The obtained results were processed using the PCS-Size mode v1.61 software.

The incubation medium consisted of 20 mM Hepes (pH 7.4, 37°C) 120 mM KCl, 2 mM potassium-phosphate buffer (pH 7.4, 37°C), 5 mM pyruvate, and 5 mM succinate.

### Detection of NADH fluorescence in mitochondria using spectrofluorometry

The relative values of NADH intrinsic fluorescence were determined with Quanta Master 40 PTI (Canada) using the FelixGX 4.1.0.3096 software. The detection was conducted in 2 mL of the following medium: 20 mM Hepes (pH 7.4 at 37°C), 2 mM K^+^-phosphate buffer (pH 7.4 at 37°C), 120 mM KCl, 5 mM pyruvate, and 5 mM succinate. The protein concentration in the sample was 50 µg/mL.

### Simulation of DCF-fluorescence, mitochondrial swelling and changes in NADH fluorescence

For the simulation, we chose the Cell Illustrator v.3 software (Human Genome Center, University of Tokyo, Japan), the basis of which is a hybrid functional Petri net. A Petri net is a directed bipartite graph with two types of nodes (Table 1): places and transitions, which are connected by arcs, reflecting the structure of the net. Places usually characterize the objects, elements, and resources of the modeled system; transitions are the events that occur in the system and the logical conditions of their implementation [17].

**Table 1.**
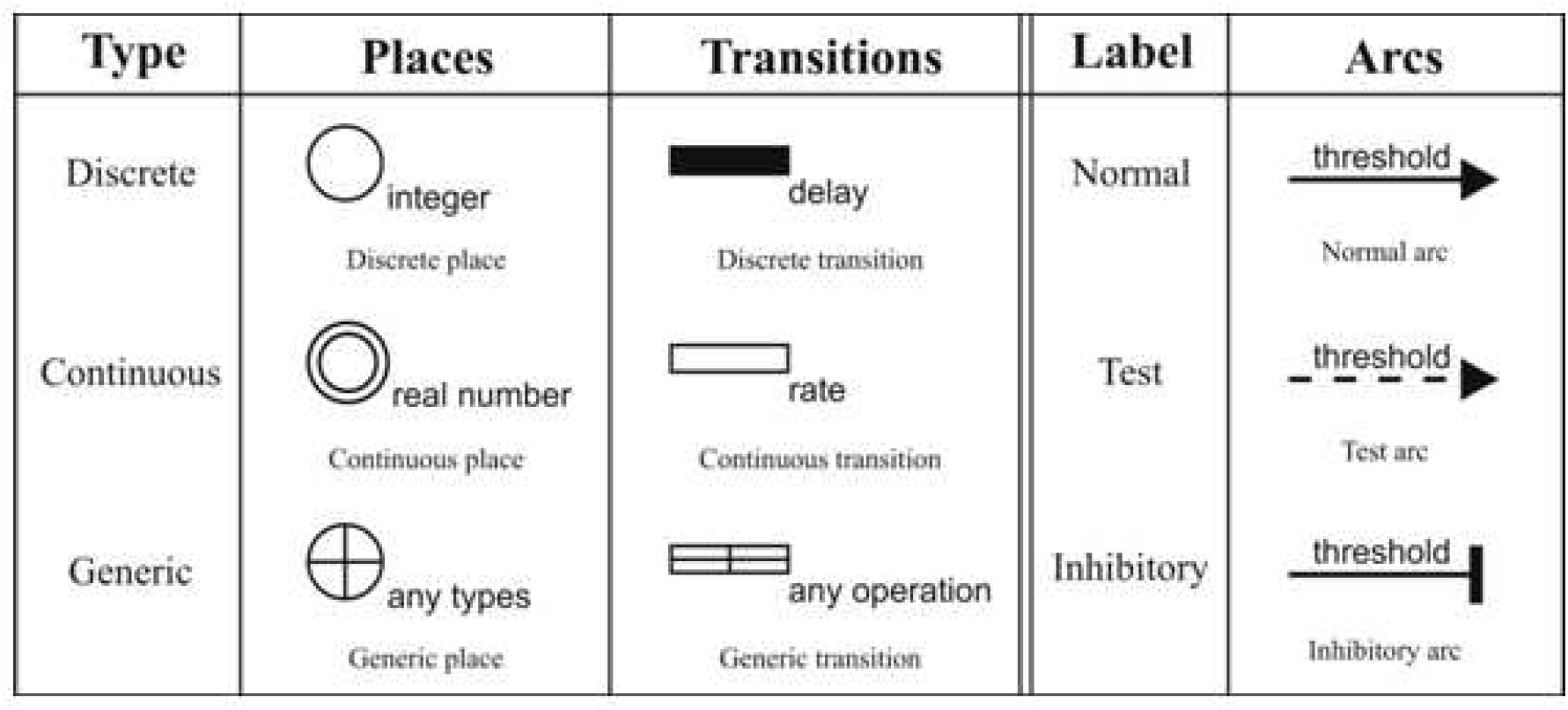
Main structural elements of the hybrid functional Petri nets (see the main text for the explanation)

| Type | Places | Transitions | Label | Arcs |
| --- | --- | --- | --- | --- |
| Discrete | integer<br>Discrete place | delay<br>Discrete transition | Normal | threshold<br>Normal arc |
| Continuous | real number<br>Continuous place | rate<br>Continuous transition | Test | threshold<br>Test arc |
| Generic | any types<br>Generic place | any operation<br>Generic transition | Inhibitory | threshold<br>Inhibitory arc |

### Assessment of the content of ionized calcium in the mitochondria

Loading of the mitochondria by the probe Fluo-4 AM at a concentration of 2 µM was carried out in the medium that contained (mM): Hepes – 10 (pH 7.4, 37°C), sucrose – 250, and 0.1% bovine serum albumin for 30 min at 37°C. To improve the process, the dye was mixed with Pluronic F-127 (0.02%). The relative values of Ca^2+^ content in the matrix of mitochondria, loaded with Fluo-4 AM (λ_ex_ = 490 nm, λ_fl_ = 520 nm) was investigated using the fluorometric method on spectrofluorometer Quanta Master PTI 40 (Canada) with software FelixGX 4.1.0.3096. The medium, from which energy-dependent accumulation of Ca^2+^ was carried out by the mitochondria, had a composition (mM): Hepes – 20 (pH 7.4, 37°C), sucrose – 250, potassium phosphate buffer – 2 (pH 7.4, 37°C), MgCl_2_ – 3, ATP – 3, sodium succinate – 5, concentration of Ca^2+^ – 80 µM [18].

***In the work the following reagents were used***: Hepes, glucose, sucrose, sodium succinate, sodium pyruvate, bovine serum albumin, ATP, NaN_3_, Pluronic F-27, DCF-DA (2’,7’-dichlorodihydrofluorescein diacetate), EGTA, CaCl_2_, A-23187 (Sigma, USA); Fluo-4 AM (Invitrogen, USA). Any other reagents are produced in Ukraine.

***The solutions were prepared*** on bidistilled water, which had a specific electrical conductivity of not more than 2.0 µcm / cm. The electrical conductivity of the water was recorded using a conductometer OK-102/1 (Hungary).

## RESULTS AND DESCUSSION

We have developed a simulation model (Fig. 1) that connects changes in the endogenous NADH fluorescence (Fig. 2), production of reactive oxygen species (Fig. 3) and mitochondria swelling (Fig. 4) for the formalization of experimental data in this work, in order to carry out the predictive function, and also to find the correspondence between the theoretical predictions and real results. We have used the "indirect donor" of NO sodium azide as biologically active compound that cause changes in the corresponding bio(physical)chemical parameters of the mitochondria (Fig. 3,4).

**Fig. 1.**
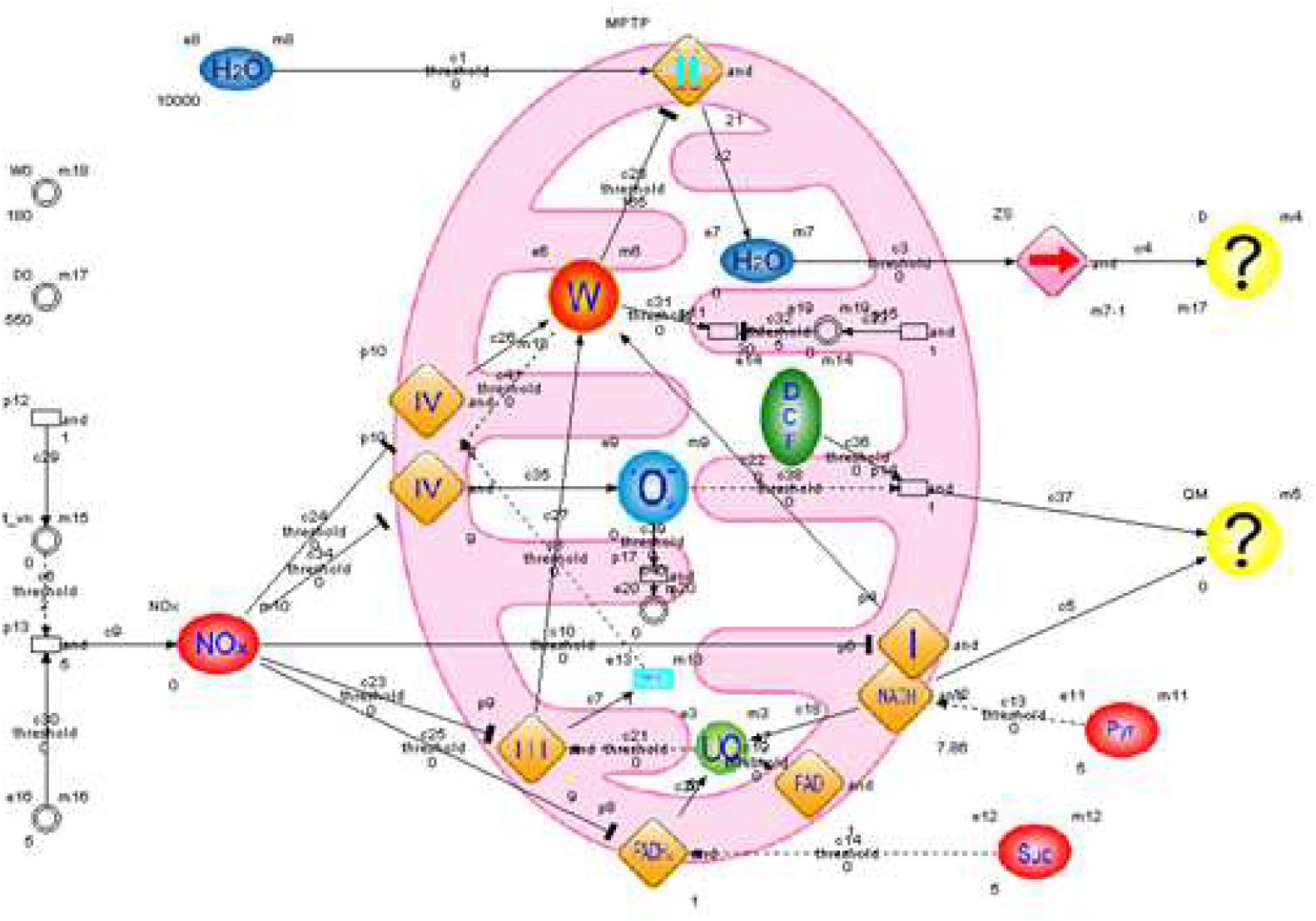
Structure of hybrid functional Petri nets, which simulated the sodium azide effect on the changes in the hydrodynamic diameter and the NADH, DCF fluorescence of the myometrium mitochondria

**Fig. 2.**
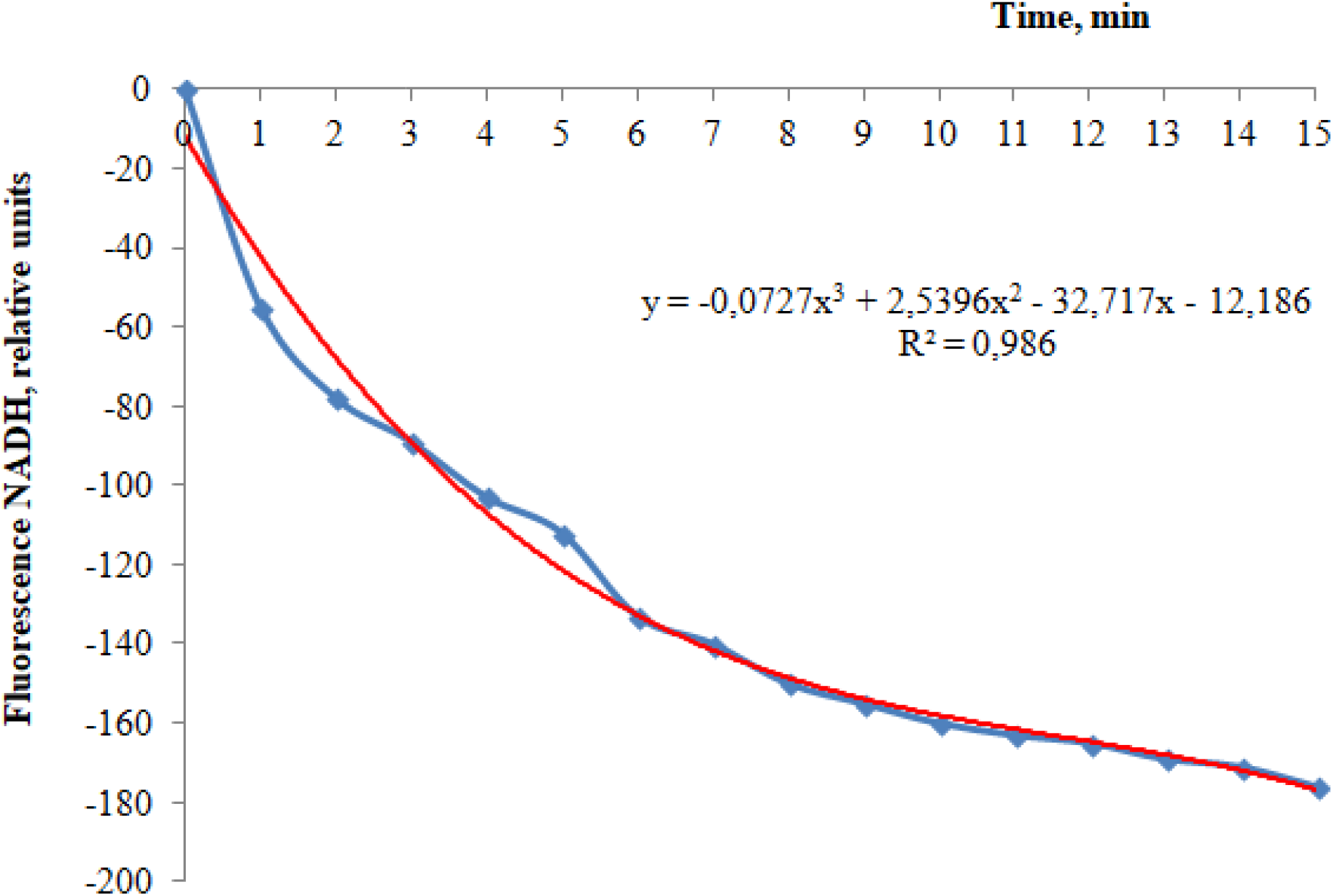
Changes in fluorescence of NADH in the isolated mitochondria from the myometrium cells. These data represent a typical experiment

**Fig. 3.**
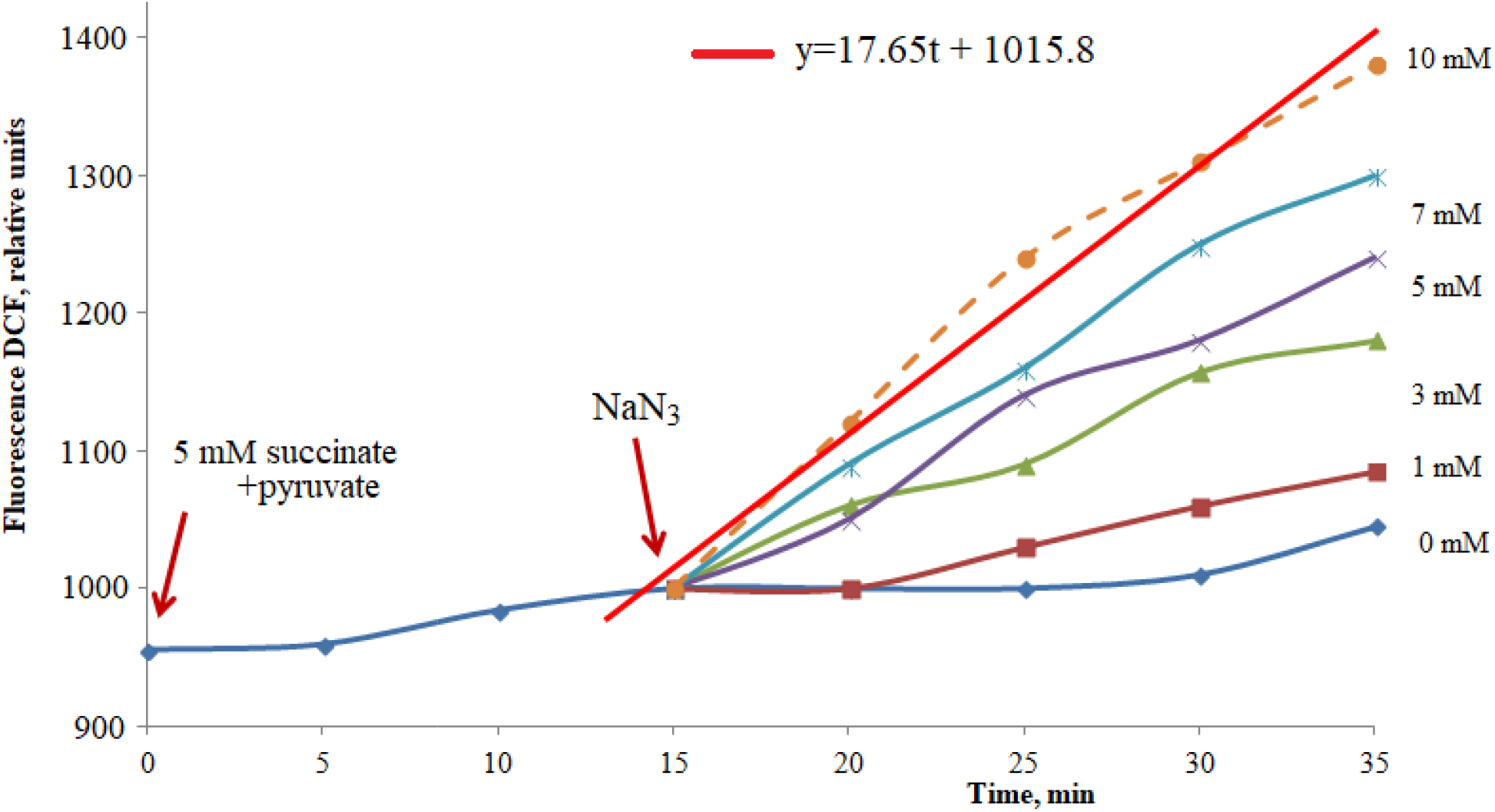
The concentration-time dependencies of the DCF fluorescence in the isolated mitochondria. The straight red line is theoretically calculated according to equation (8) for 10 mM sodium azide; dotted brawn line – experimental obtained curve for 10 mM sodium azide. These data represent a typical experiment

**Fig. 4.**
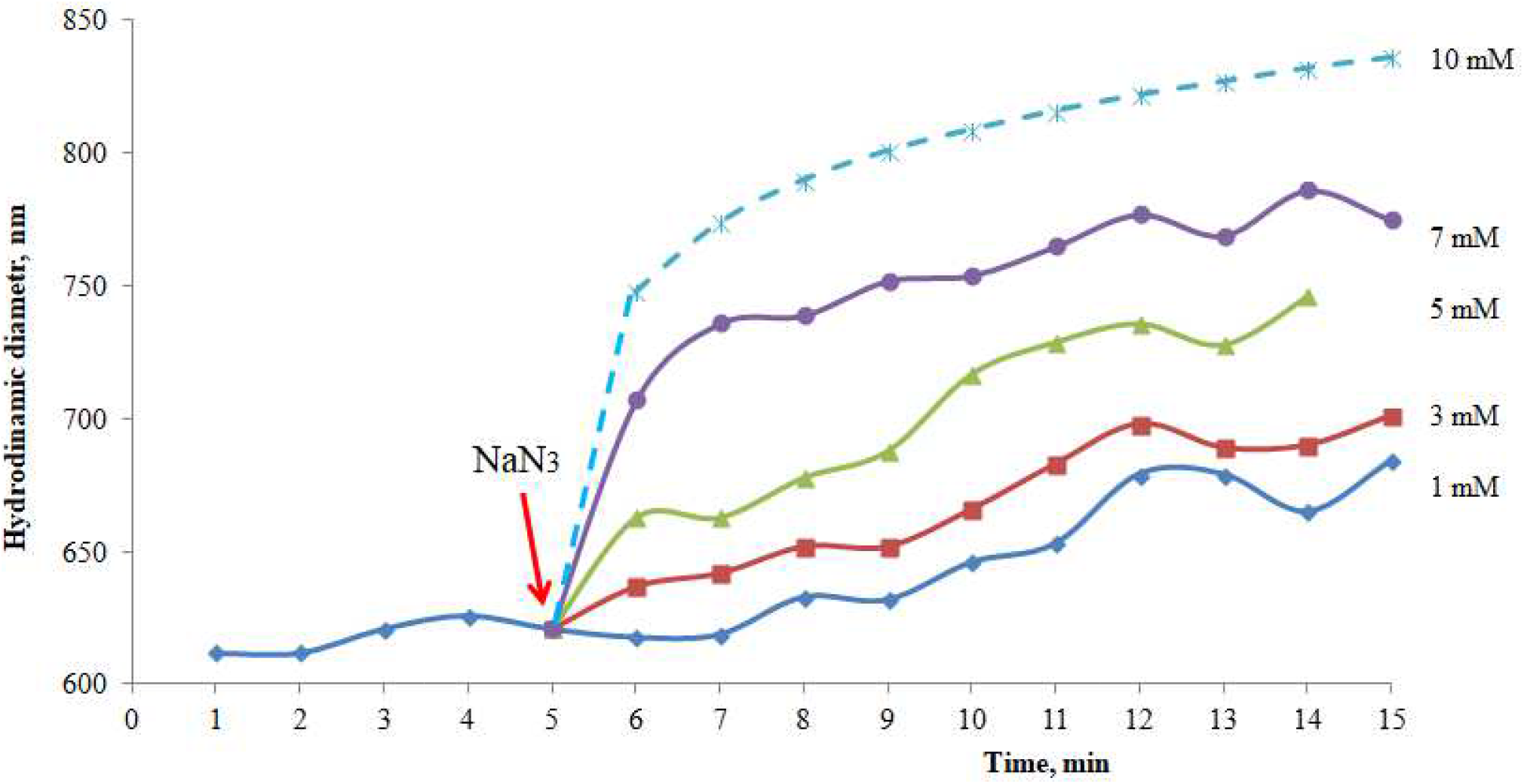
The values of mitochondria hydrodynamic diameter during swelling on NaN_3_ action. The curve for 10 mM sodium azide is theoretically calculated according to equation (9). These data represent a typical experiment

It was established, that NaN_3_ increased the mitochondria hydrodynamic diameter (causing swelling) and amplified their DCF fluorescence (it stimulated the formation of reactive oxygen forms), depending from its concentration (1-3-5-7 mM) and time (0-15 min), see Fig. 3, 4. Inhibition of the respiratory chain *IV* complex results to the slowing down of the mitochondria functioning, the reduction of the electrical and chemical potential on the internal membrane (∆pH), the opening of the permeability transient pore (MPTP) and the mitochondria swelling [19–23]. The accumulation in the electron transport chain of the semi-reduction components, especially in Q-cycle, leads to an increase in the process of one-electron oxygen reduction [24]. Thus, the generation of superoxide anion and other reactive oxygen species will be intensified. Simultaneously, the chemical decomposition of sodium azide with the formation of nitrogen oxides will be accompanied by an increase in the generation of peroxynitrite, which contributes to DCF fluorescence too. Strengthening under these conditions the content of reactive nitrogen and oxygen species in mitochondria will further reduce the functional activity of the electron transport chain and matrix enzymes, especially those containing iron-sulfur centers and metal proteins [25–27]. The latter will result in the processes of the mitochondrial dysfunction, collapse of bioenergetics up to cell death [21,23,28]. Using Petri hybrid functional nets, we simulated the effect of sodium azide, depending on the time and concentration, on the hydrodynamic diameter and DCF fluorescence of the myometrium isolated mitochondria.

The components of the mitochondria incubation medium and the effects of sodium azide on their bio(physical)chemical parameters were taken into account in the simulation. We operated the following experimental facts in the process of the model creation: (1) - succinate and pyruvate are added to the mitochondria incubation medium for their energized; (2) - sodium azide, as an "indirect NO donor ", has an inhibitory effect on the electronic transport chain as a whole, but the greatest effect on the morpho-functional parameters of the organelles has inhibition of the *I*, *III* and *IV* complexes (3) - inhibition of the electron transport chain activity results in the intensification of the reactive oxygen forms production in the mitochondria and, consequently, DCF fluorescence (direct inhibition of the *IV* complex within the framework of the simulation model); (4) - inhibition of the electron transport chain by sodium azide leads to an increase in the hydrodynamic diameter of the mitochondria: the osmotic balance between the matrix and the non-mitochondrial medium is broken due to the activation of the permeability transition pore with the subsequent H_2_O entrance to the matrix and swelling of the mitochondria (this is the result of inhibition of *I*, *III* and *IV* complexes and depolarization of the internal mitochondrial membrane in the framework of the simulation model).

Symbols in the scheme: NO_x_—nitrocompounds, particularly sodium azide; Suc— sodium succinate; Pyr—sodium piruvate; W—electric potential of the inner mitochondrial membrane; I, II, III, and IV as well as FADH_2_, FAD, and NADH (inscribed in a rhombus in the mitochondrial membrane) - complexes of the electron transport chain and the appropriate cofactors; O_2_^•-^ - superoxide anion; DCF - DCF-DA fluorescence probe; UQ—ubiquinone; cyt C—cytochrome C; symbol **-| |-** inside a rhombus—cyclosporine-sensitive permeability transition pore, MPTP; and arrows: → — activation of the process and ┬ — inhibition of the process

Structure of hybrid functional Petri nets, which simulated the sodium azide effect on the changes in the hydrodynamic diameter, NADH and DCF fluorescence of the myometrium mitochondria is presented on scheme (Fig. 1). Symbols in the scheme are explained on the caption. Some particular specifications are listed below. The place m15 is a timer that controls the time of the inhibitor insertions in the amount determined by the place m16. Before the insertion of a specific inhibitor of electron transport chain, the mitochondrial hydrodynamic diameter, measured by Zeta-sizer (transition ZS and place m4) practically does not change. The fluorescent response of NADH/DCF is registered by the spectrofluorimeter QM (place m5) and corresponds to the “control” curves (Fig. 2, 3).

The insertion of sodium azide (NO_x_, Fig. 1) breaks the work of electron transport chain that decreases the electric potential of the inner mitochondrial membrane W (place m6). It causes an activation of the mitochondrial permeability transition pore (MPTP), H_2_O transport into the matrix (place m7), and an increase in the mitochondrial hydrodynamic diameter. The fluorescence response changes are simulated by the NADH and DCF transitions with velocities in accordance with equations (5, 7) as described below.

Through the modeling, we obtained mathematical equations that formalized the process of mitochondria swelling and the changes in the NADH/DCF fluorescence in the medium supplemented with sodium azide. In particular, these equations adequately described the time characteristics of the mitochondria swelling (Fig. 4), changing of the NADH/DCF fluorescence (Fig. 2, 3) simultaneously. The permeability of MPTP and the intensity of the NADH/DCF fluorescent response are the time derivatives from the corresponding dependencies.

The substrate of pyruvate dehydrogenase complex (5 mM pyruvate) that produced NADH for electron transport chain, and 5 mM succinate, a substrate of FAD-dependent succinate dehydrogenase was into incubation medium in order to produce the energized state of mitochondria [29]. It has been shown that NADH fluorescence decreased in time in the presence of respiratory substrates, which indicates increase in NAD^+^ content resulting from functioning of NADH-ubiquinone oxidoreductase of electron transport chain (Fig. 1). We have used experimental results from reduce the auto-fluorescence of NADH in isolated mitochondria over time under the condition of the electron transport chain functioning for modeling in this study. The time dependence of the average change in the fluorescent response of NADH (autofluorescence) was approximated by polynomials as follows:

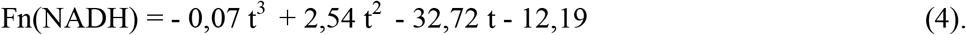

The intensity of the NADH fluorescent response:

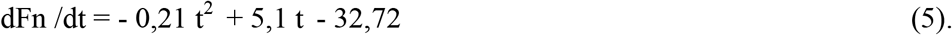

The concentration-time dependencies of the DCCF fluorescence can be approximated by polynomials as:

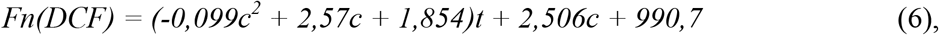

where c – concentration of sodium azide, mM, t – time, min.

The intensity of the DCF fluorescence:

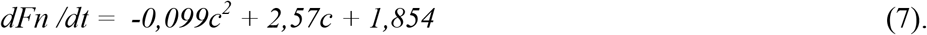

The obtained expressions make it possible to predict the time ddependence of DCF fluorescence, for example, for 10 mM concentration of sodium azide (Fig. 3):

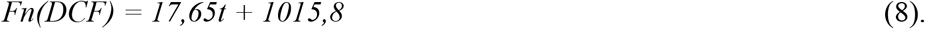

According to results for 1-7 mM NaN_3_ (Fig. 4) dynamics of the concentration-dependent changes in the average values of mitochondria hydrodynamic diameter during swelling can be approximated as:

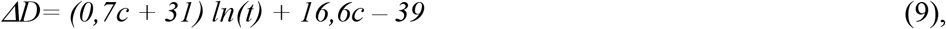

where c – concentration of sodium azide, mM, t – time, min.

So, the permeability oof mPTP is described as:

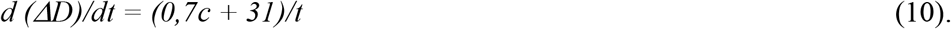

It is possible to predict theoretically the course of the corresponding curves for unknown sodium azide concentrations based on this equation. Fig. 4 shows an example of such a curve for 10 mM sodium azide.

Our model enabled a simultaneous prediction of the changes in the organelle NADH/DCF fluorescence and their hydrodynamic diameter in time, which enabled us to significantly optimize the time of the experimental procedures, the consumption of reagents, and the use of laboratory animals. Moreover, it allowed us to analyze the dynamics of the processes and to compare the modeling results with the actual observations, considering the changes in the abovementioned parameters (the composition of the incubation medium and the presence of the activators/inhibitors).

Sodium azide is known to degrade in water solutions, producing hydrazoic acid, hydroxylamine, and, possibly, nitrogen oxides, which act as reactive nitrogen species in biological systems [12]. In our previous experiments sodium azide (5 mM) caused more pronounced decrease in NADH fluorescence than in control, and also increase in FADH_2_ contents, which may indicate blocking of electron transfer from succinate to ubiquinone [30]. We have attributed the decreased NADH and FAD levels under nitrocompounds to drop in activity of enzymes of citric acid cycle due to inhibition of electron transport chain in mitochondrial membrane. Thus, we investigated with flow cytometry the effect of the sodium azide on matrix content of Ca ions in mitochondria to confirm the effect of NO_x_ on the mitochondria functional activity (Fig. 5). Addition of exogenous Ca^2+^ to mitochondria suspension was associated with the increase in fluorescence of Fluo-4 that had been loaded into them in advance, which indicates increased matrix Ca^2+^ concentration. The cation was accumulated in the presence of Mg-ATP^2^- and succinate for 5 min, at which moment the stable level of Ca^2+^ accumulation was achieved (Fig 5, column 1). We ascertained the barrier function of mitochondrial inner membrane towards Ca ions by addition of A23187 Ca^2+^-ionophore to suspension. This was associated with rapid release of the accumulated cation (Fig 5, column 2). The sodium azide caused efficient release of accumulated Ca^2+^ from mitochondria (Fig 5, column 3). The deenergizing effect of sodium azide on mitochondria has been reliable established. The slight increase in ionized Ca in the matrix was observed, as well as a significant decrease in Ca accumulation in comparison with the control, under conditions of mitochondrial pre-incubation with sodium azide (depolarization of the internal membrane) (Fig. 5, column 4 and 5 respectively).

**Fig. 5.**
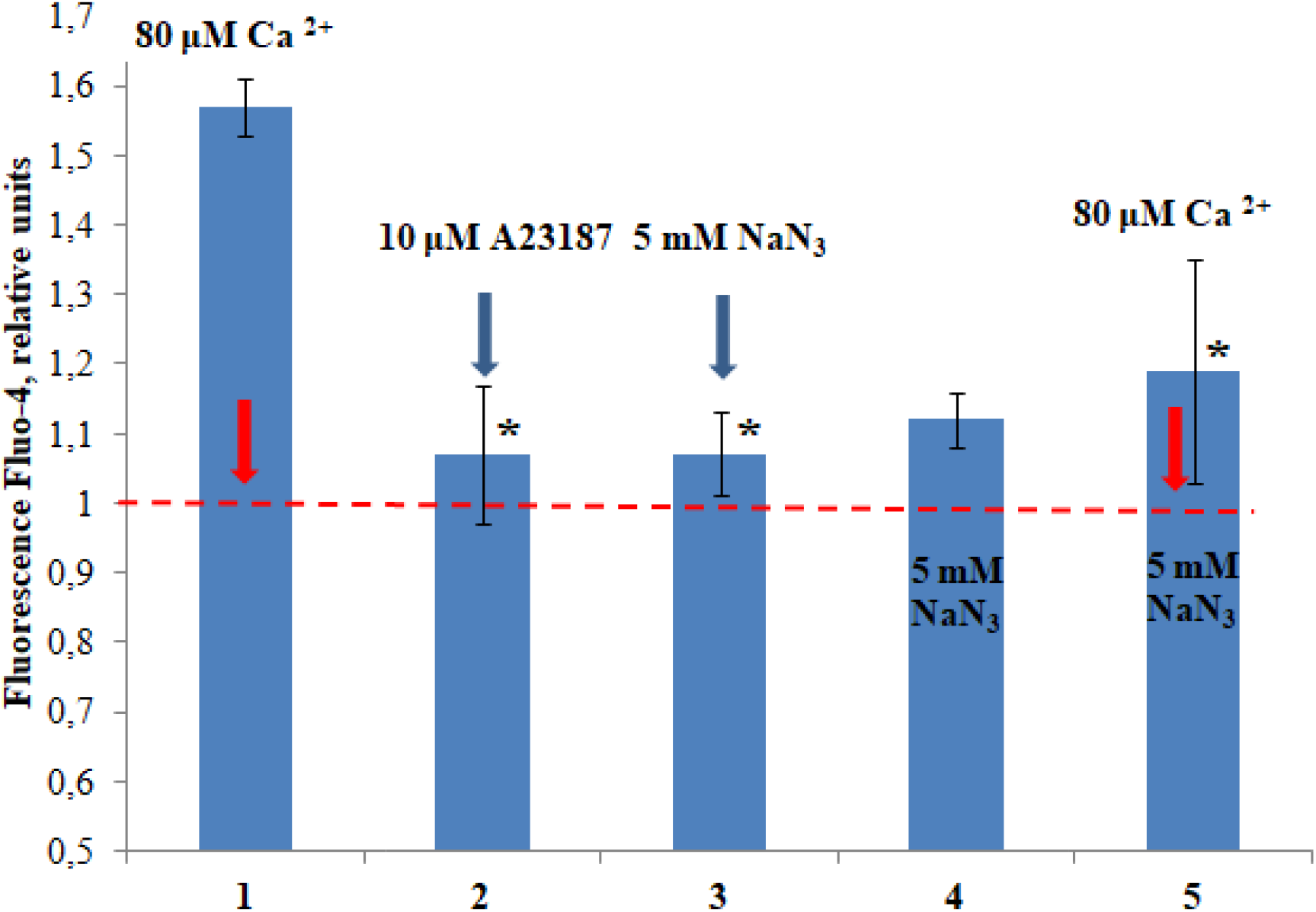
Effects of Ca^2+^-ionophore and sodium azide on matrix Ca^2+^ content. The cation had been accumulated in energy-dependent manner. Endogenous ionized Ca – “1”, starting level. The data are presented as mean ± SEM, * - P ≤ 0.05, n=5

Thus, using the Petri fluorescence of endogenous nets, the simulation model for simultaneous changes in the NADH, the characteristic size of mitochondria, and the generation of reactive oxygen species under real experimental conditions has been created, which combines functional changes with the structural representation of these processes. The simulation results in mathematical equations that formalize simultaneous processes of mitochondrial swelling and DCF fluorescence in a medium with NaN_3_. The de-energizing effect of sodium azide on mitochondria results in disturbance in their functioning, namely the ability to effectively accumulate and maintain Ca^2+^ in their matrix.

## CONCLUSIONS

The simulation results in mathematical equations that formalize simultaneous processes of mitochondrial swelling, changes in NADH and DCF fluorescence in a medium in which NaN_3_ is presented. These equations are able to adequately describe the time and concentration characteristics of these processes, as well as predict the intensity of their occurrence. In particular, the response of mitochondria over time to the action of NaN_3_ in a concentration, that was not used in the experiment, was predicted. The calculated values of the studied bio(physical)chemical parameters were consistent with experimentally obtained data. Creation of an adequate model optimizes experimental procedures (time, costs of reagents and laboratory animals), allows analyzing the dynamics of processes and comparing experimental results with theoretical calculations provided that the composition of the incubation medium changes.

